# An Oatp transporter-mediated steroid sink promotes tumor-induced cachexia in *Drosophila*

**DOI:** 10.1101/2020.11.05.370411

**Authors:** Paula Santabárbara-Ruiz, Pierre Léopold

## Abstract

Cancer cachexia is a multifactorial syndrome associated with many types of tumors and characterized by a combination of anorexia, loss of body weight, catabolic alterations and systemic inflammation. We developed a tumor model in *Drosophila* larvae causing a cachexia-like syndrome, and used it to evaluate the role of steroid hormone imbalance in cachectic alterations. Cachectic larvae show reduced levels of the circulating steroid ecdysone. Artificially importing ecdysone in the tumor using the Oatp74D importer aggravates cachexia, while feeding animals with ecdysone rescues cachectic defects. This suggests that a steroid sink induced by the tumor promotes catabolic alterations in healthy tissues. We find that Oatp33Eb, another member of this family of transporters, is specifically induced in tumors promoting cachexia. Blocking Oatp33Eb in cachectic tumors restores circulating ecdysone and reverses cachectic alterations. Oatp transporters are induced in several types of hormone-dependent tumors, suggesting that a similar sink effect could modify hormonal balance in cachectic cancer patients.

**Highlights:**

- We used the *Drosophila* larva as a model to study hormonal imbalance in tumor-induced cachexia.
- Steroid (ecdysone) levels are reduced in cachectic tumor-bearing larvae.
- Steroid import by the tumor reduces circulating hormone levels and induces cachexia.
- The Oatp33Eb transporter is specifically upregulated in cachectic tumors and its inhibition rescues both circulating steroid levels and metabolic/tissue homeostasis.
- Specific *Oatp* genes are found upregulated in human tumors with high prevalence of cachexia.

## Introduction

Cachexia is a devastating tumor-induced metabolic disorder marked by a progressive wasting of adipose and muscle tissues. This is driven by a combination of reduced food intake, insulin resistance, excess catabolism and inflammation. Cachexia greatly impacts the quality of life of patients, reduces the efficacy of chemotherapy and increases susceptibility to infections. This condition is often poorly diagnosed and very few opportunities for treatment are offered (Fearon, Arends and Baracos, 2013).

Rodent models have provided important contributions to our understanding of cancer cachexia (Bennani-Baiti and Walsh, 2011; Molfino *et al*., 2019). They established a role for two major groups of pro-catabolic agents: (i) tumor-produced factors (pro-inflammatory cytokines, proteolysis-inducing factors and lipid mobilizing factors) and (ii) host factors, mainly pro-inflammatory cytokines.

In parallel, patient studies have highlighted an important hormonal imbalance associated with the cachectic syndrome. Cachectic patients usually present high GH and low IGF-I levels, suggesting a state of GH resistance (Honors and Kinzig, 2012). Insulin resistance is also frequently observed in these patients. Finally, hypogonadism is found associated with cachexia in men and lower circulating testosterone correlates with reduced muscular mass. The increase in catabolic processes suggests that cachexia could at least partially result from a reduction of general anabolic factors, and motivated the use of hormone replacement therapies. However, the molecular mechanisms responsible for these hormonal perturbations remain to be elucidated, and are needed for novel therapeutic approaches.

Several *Drosophila* tumor models have recently been used to study cancer-induced cachexia, taking advantage of the conservation with vertebrates of both the pathways regulating tumor growth and the hormonal physiology (Brumby and Richardson, 2005; Leopold and Perrimon, 2007; Droujinine and Perrimon, 2016). The main advantages of these genetically-engineered fly models is that the tumor and the host can be modified in parallel, giving the opportunity to study intricate tumor-host interactions.

In the adult fly, several tumor-produced factors have been defined as cachectic inducers, such as the IGF-binding protein (IGFBP) homolog ImpL2 and the PDGF/VEGF homolog Pvr1 (Figueroa-Clarevega and Bilder, 2015; Kwon *et al*., 2015; Song *et al*., 2019). However, the adult fly model presents some limitations due to the technical difficulty to perform tumor injections avoiding an immune challenge and controlling the quantity of injected tissue, or the possible bias of generating tumors in organs essential for animal physiology.

An alternative is to observe metabolic alterations in larvae developing epithelial tumors in non-essential tissues such as the wing or eye imaginal discs. Larval physiology relies on conserved hormonal relays like insulin-like peptides, glucagon-like hormones and steroids (ecdysone), which make it a particularly suited system to study how tumor development impairs general metabolic homeostasis and hormonal balance.

In the current work, we use a larval model of tumor-induced cachexia to study the role of hormone imbalance in this process. Cachectic, tumor-bearing animals show an important reduction of circulating steroids (ecdysone) and a state of insulin resistance, suggesting a conserved hormonal imbalance. Artificially inducing steroid import in tumor cells by expressing the ecdysteroid transporter Oatp74D (also called Ec-I) aggravates the proportion of cachectic animals without increasing tumor burden. Conversely, feeding cachectic animals with ecdysone-containing food rescues cachexia. Oatp33Eb, another member of the Oatp transporters (and not other family members), is specifically induced in cachectic tumors, and the knock-down of this gene in the tumor restores normal levels of circulating ecdysone and rescues all cachectic symptoms, while maintaining tumor growth.

We therefore propose a model whereby tumors induce cachectic symptoms by uptaking circulating steroids at the expense of healthy tissues. This “sink effect” leads to a catabolic switch as observed in muscles and fat. Remarkably, some members of the OATP family of transporters are induced in human tumors prone to cachectic evolution, opening the possibility to use this mechanism of steroid uptake as an early marker or a therapeutic target of cancer-induced cachexia.

## Results

### Tumor-induced cachexia triggers hormonal imbalance in *Drosophila* larvae

To explore *in vivo* the role of hormonal imbalance in tumor-induced cachexia, we used *Drosophila* larvae as a fast and reproducible model that recapitulated several of the pathophysiological mechanisms observed in the human syndrome. To this aim, we induced tumor development in the eye imaginal disc, a non-essential tissue with no described physiological function during larval life, using the Flip-out technique (Pagliarini and Xu, 2003 - see methods). The combination of an oncogenic *ras*^*V12*^ mutation with an interference RNA silencing the tumor-suppressor gene *disc large* (*dlg*^*RNAi*^) induces aggressive tumors. After 3 days these tumors drive cachectic symptoms in a fraction of the larval population. Cachectic larvae present an accumulation of body fluid (bloating syndrome), body weight loss, fat and muscle tissue wasting, and a perturbation of feeding behavior (anorexia) (Fig. 1A-G, Suppl. Fig. 1A-E). In addition, we observed that the fat body (the functional equivalent to the mammalian fat and liver) of cachectic animals responds poorly to insulin stimulation (Fig. 1H, Suppl. Fig. 1F). This, together with high blood glucose, insulinemia and reduced glucose uptake by fat body cells (Fig. 1I, Suppl. >Fig. 1F,G), defines a state of insulin resistance, as described in cancer cachexia. Notably, Impl2, a recently described cachectic inducer interfering with insulin response was upregulated in tumors, but its inhibition did not rescue cachexia (Suppl. Fig. 2C, G).

**Figure 1.**
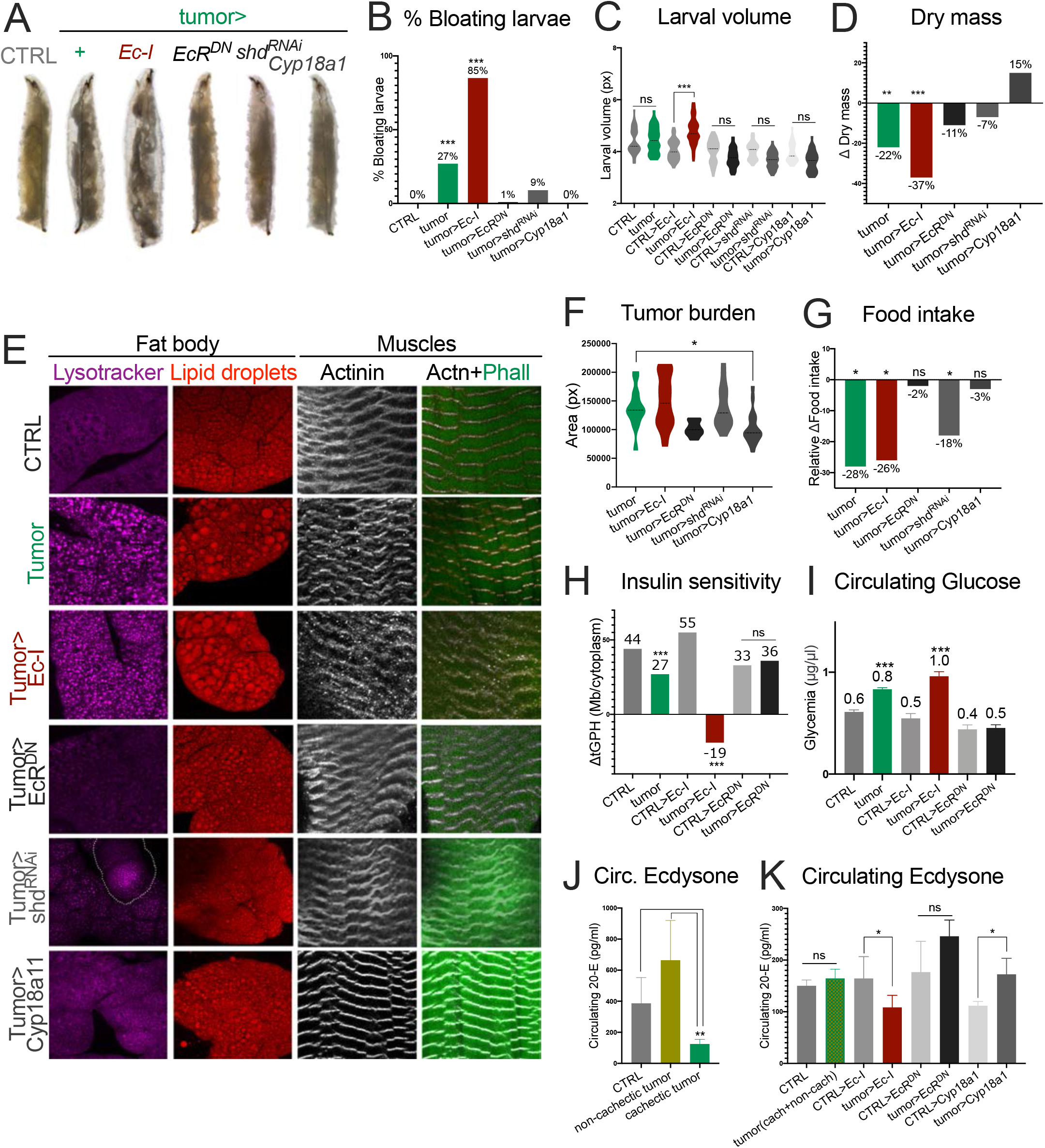
Tumor-induced cachexia triggers hormonal imbalance in *Drosophila* larvae. (A,B) Representative images of control and tumor-bearing larvae showing bloating (A) and percentage of bloating animals with the indicated genotypes (n=200 for each genotype). Only *tumors>+* and *tumors>Ec-I*-larvae showed bloating (27% and 85%, respectively). ***p<0,001. (C) Quantification of larval volumes with the indicated genotypes (n=50 for each genotype). ***p<0,001. (D) Differences in dry mass (calculated with the formula dry mass*100/wet mass) in larvae with the indicated genotypes (n=50 for each genotype). In each column, the delta between *CTRL* and tumor conditions is plotted. Bloated *tumor>* and *tumors>Ec-I* larvae show significant dry weight loss. **p<0,05, ***p<0,001. (E) Peripheral tissue wasting in tumor-bearing larvae. Lysotracker labels acidic vesicles (purple) and Nile Red labels lipid droplets (red) in fat body cells from larvae with the indicated genotypes. αActinin (white) and Phalloidin (green) stainings show larval muscle morphology. The dotted line surrounds a GFP positive cell in *tumors>shd*^*RNAi*^. (F) Tumor size in larvae with the indicated genotypes (n=30 per genotype). The dotted line represents the mean. *p<0,05. (G) Differences in food intake measured using blue dye Erioglaucine in the food (n=12 groups of 8 larvae each). Values relatives to tumor-less controls are showed, *p<0,05. (H) Insulin sensitivity in fat body cells by measuring the delta in membrane *tGPH* in fat bodies incubated with or without insulin. The delta for Ctrl and various tumor conditions is plotted. ***p<0,001. See Suppl. Fig. 1F for confocal images. (I) Concentration of hemolymph glucose from larvae with the indicated genotypes (mean +/-SEM, ***p<0,001). The plotted values correspond to the mean. (J,K) Quantification of circulating 20-E (pg/ml/well) found in the hemolymph of larvae with the indicated genotypes (mean +/-SEM, *p<0,05). Note that in J, tumor-bearing animals were divided into non-cachectic and cachectic subgroups, whereas in K, cachectic and non-cachectic are plotted together in the tumor group.

**Figure 2.**
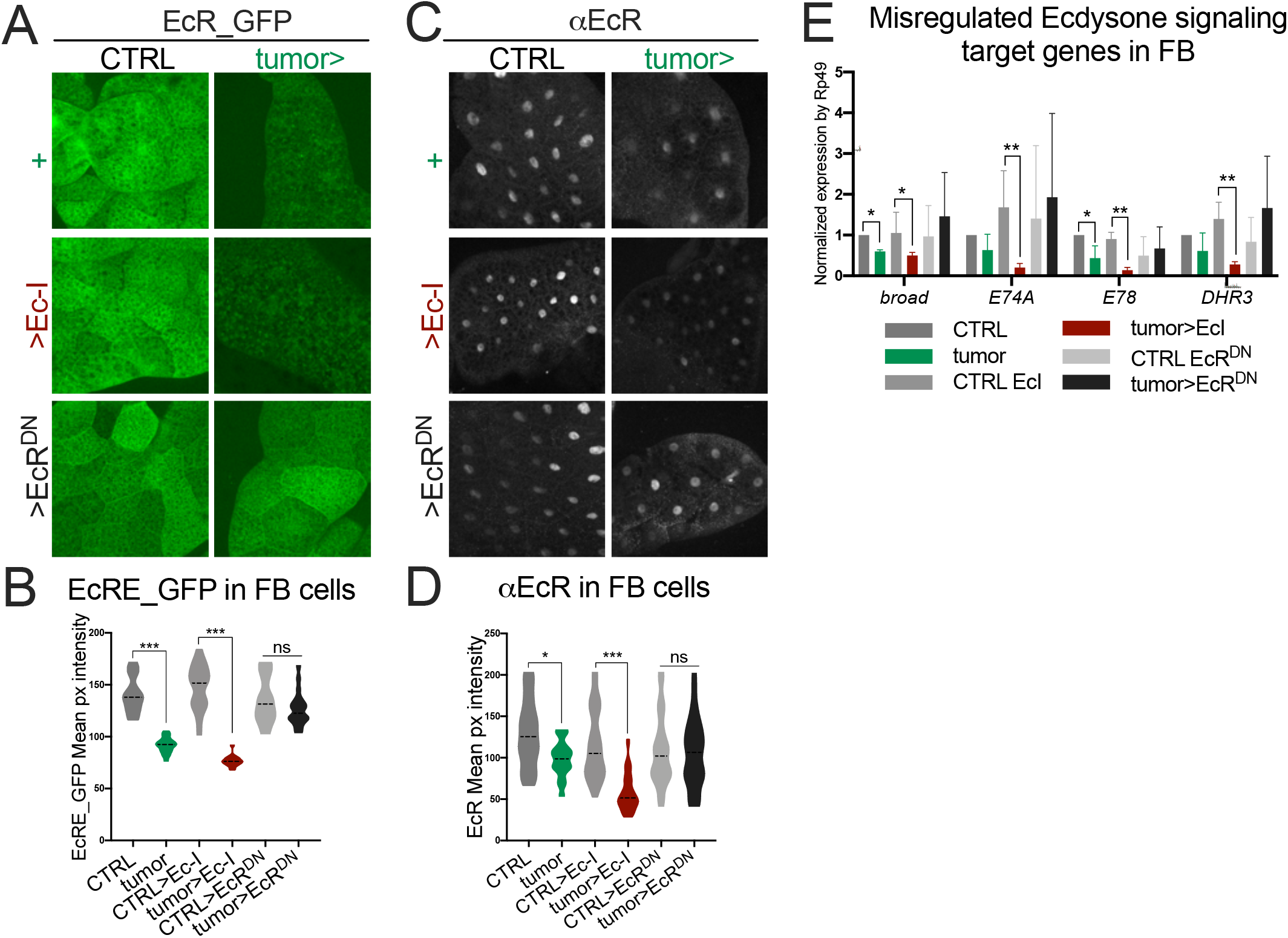
Ecdysone uptake by the tumor causes a decrease in ecdysone signaling in healthy tissues (I). (A,B) (A,B) Fat bodies from larvae with the indicated genotype carrying the EcRE-GFP reporter stained with αGFP (green) and quantification of the EcRE-GFP pixel intensity (B). The dotted line represents the mean. ***p<0,001. (C,D) EcR staining in fat body cells from animals with the indicated genotype (C, white) and quantification of the αEcR pixel intensity (D). The dotted line represents the mean. *p<0,05, ***p<0,001. (E) mRNA levels of ecdysone targets (*broad, E74A, E78* and *DHR3*) in fat bodies from different genotypes, measured by RT-PCR. The results shown are mean +/-SEM. *p<0,05, **p<0,01.

The steroid hormone ecdysone (E) plays key roles during *Drosophila* development both as an anabolic hormone and a timer of developmental transitions (Yamanaka, Rewitz and O’Connor, 2013). We first noticed that tumor-bearing cachectic larvae present reduced circulating levels of ecdysone, compared to tumor-bearing, non-cachectic animals (Fig. 1J). These animals present an upregulation of Dilp8 and a 6hr delay at the larva-pupa transition (Suppl. Fig. 2A,B). Since Dilp8 could inhibit ecdysone production, we tested the effect of silencing Dilp8 in the tumor on the developmental delay and the cachectic behavior. In these conditions, although the developmental delay was rescued, the prevalence of cachexia was slightly modified (Suppl. Fig. 2A,C). Furthermore, we did not observe differences in the levels of expression of the “Halloween” group of ecdysteroidogenic enzymes in cachectic animals, suggesting that ecdysone synthesis is not modified by the presence of the tumor (Suppl. Fig. 2D). This was confirmed by the observation that the PTTH and Serotonin circuitries were intact in the tumor-bearing larvae (Suppl. Fig. 2E). Therefore, the reduction of E levels in cachectic larvae is due to another mechanism. To test this hypothesis, we forced ecdysone uptake in the tumor by inducing the expression of Ec-I, a member of the SLCO transporter family recently shown to specifically import ecdysone in *Drosophila* cells (also called Oatp74D, Okamoto *et al*., 2018). Although neither tumor burden nor aggressiveness were significantly modified in these conditions, the percentage of cachectic animals increased from 27% to 85% (compare *tumor>* and *tumor>EcI*, Fig. 1A-I, Suppl. Fig. 1A-G, Suppl. Fig. 2F), correlating with a reduction of circulating ecdysone in the general population of tumor-bearing larvae (Fig. 1K). Conversely, blocking ecdysone signaling in the tumor (*tumor>EcR*^*DN*^, *tumor>shd*^*RNAi*^ and *tumor>Cyp18a1*, see methods) rescued circulating levels of ecdysone, as well as cachexia, suggesting that a positive feedback mechanism involving intra-cellular EcR signaling controls ecdysone uptake by the tumor (Fig. 1A-I, Suppl. Fig. 1A-G, Suppl. Fig. 2F).

In conclusion, a fraction of tumor-bearing larvae presents cachectic symptoms associated with hormonal imbalance including a reduction of circulating steroids. Increasing ecdysone uptake by the tumor reduces its circulating levels and aggravates the cachectic effect without affecting tumor growth or aggressiveness. This suggests that the catabolic switch observed in non-tumorous tissues could be due to an ecdysone “sink effect” induced by the tumor.

### Ecdysone uptake by the tumor causes a decrease in ecdysone signaling in healthy tissues

The consequence of a sink effect would be a reduction of ecdysone signaling in peripheral tissues. To test this, we quantified Ecdysone receptor (EcR) activity using the reporter EcRE-GFP (Kimura *et al*., 2008) in fat body cells. Indeed, the level of GFP fluorescence in the fat of tumor-bearing animals (*tumor>*) is reduced compared to non tumor-bearing ones (Fig. 2A, B). The quantification shows a further decrease of EcRE-GFP in *tumor>EcI*, and a rescue in *tumor>EcR*^*DN*^ (Fig. 2A, B). We also observed a significant reduction of nuclear EcR labeling in the fat body of cachectic animals (Fig. 2C, D). This was confirmed by analyzing the level of direct targets of EcR like *broad* and *E78*, which show a reduction in *tumor>* and *tumor>EcI* conditions (Fig. 2E). This indicates that variations in circulating E correlate with the level of EcR signaling in peripheral tissues, and that the uptake of E by the tumor limits ecdysone signaling in peripheral tissues.

We then tested whether supplementing ecdysone in the food would rescue the cachectic phenotype. Indeed, adding 20-E, the active form of E, in the fly food efficiently rescues the cachectic fraction both in *tumor>* and in *tumor>EcI* conditions without modifying the tumor burden or aggressiveness (Fig. 3A-G, Suppl. Fig. 3). Conversely, feeding animals with a food medium lacking ergosterol (a precursor of ecdysone that is no longer produced by the *erg2?* yeast strain) strongly increased the proportion of cachectic animals (27% to 100%, 120hr AED) despite a reduced tumor burden (Fig. 3H-O, Suppl. Fig. 4). Remarkably, although *tumor>EcR*^*DN*^ animals did not show cachexia in normal food, they all presented important bloating in *erg2?* food (Fig. 3H-O, Suppl. Fig. 4). This indicates that the reduction of circulating steroid is the main trigger for inducing cachexia in tumor-bearing larvae.

**Figure 3.**
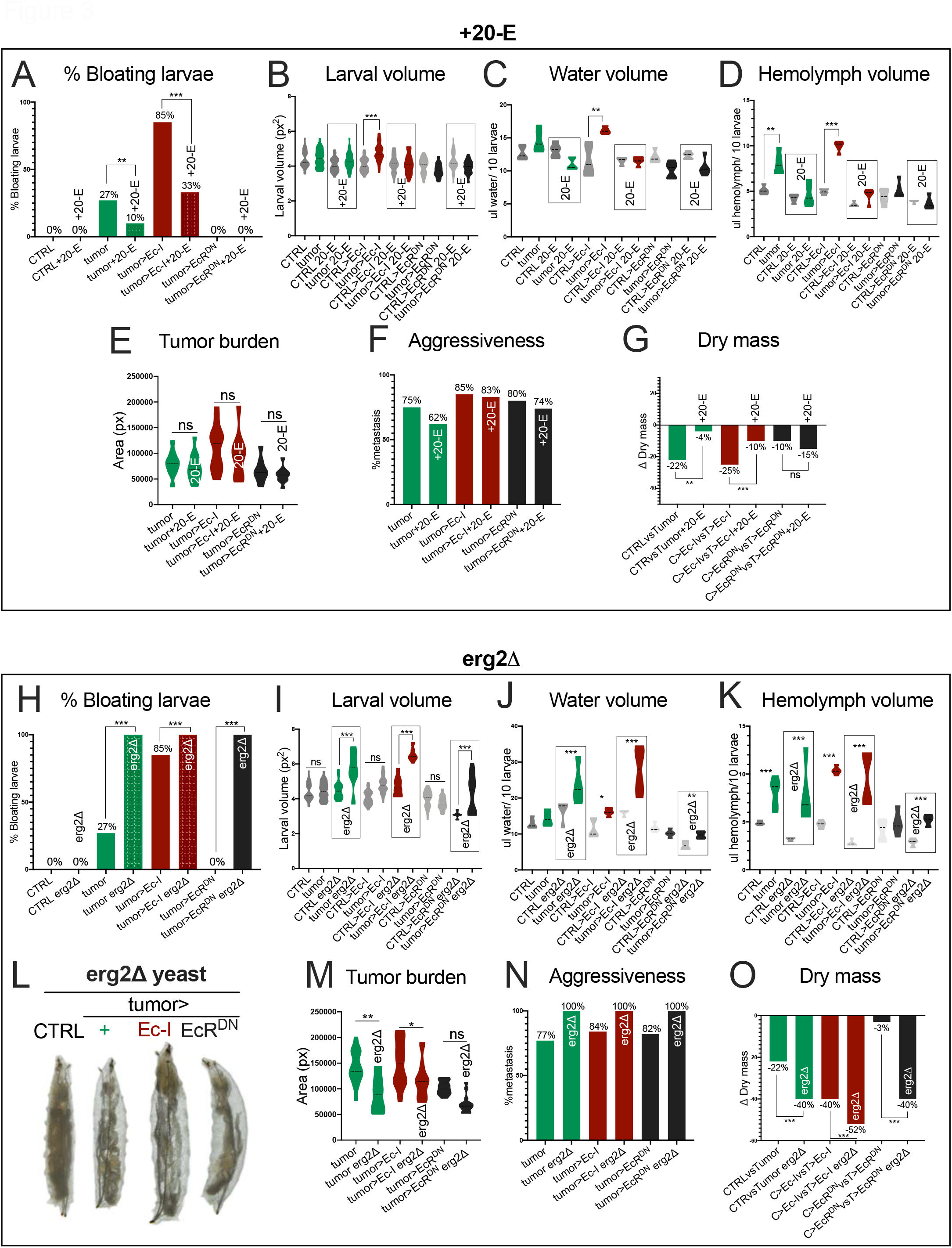
Cachexia inversely correlates with the levels of circulating Ecdysone. (A-D) The cachectic phenotype is rescued by the addition of 20-E. Percentage of animals with bloating defects (A, n=200 for each genotype), mean larval volume (B, n=50 for each genotype), water volume (C, n=50 for each genotype), and extractable hemolymph (D, n=50 for each genotype) from larvae reared on standard food versus food containing exogenous 20-E, with the indicated genotypes. dotted line represents the mean. **p<0,01, ***p<0,001. (E,F) Tumor sizes (E, n=30 per genotype) and metastasis (F, n=200 for each genotype) in larvae reared on standard food versus food containing 20-E. In F, the dotted line represents the mean, no significant differences were found. (G) Differences in the content of dry mass observed in larvae with the indicated genotypes fed with standard vs 20-E-food (n=50 for each genotype). The delta between CTRL and tumor-bearing animals is plotted as a single value. ***p<0,001. (H-K) The cachectic phenotype is aggravated on *erg2Δ* yeast-containing food. Percentage of animals with bloating defects (H, n=40 for each genotype), larval volume (I, n=40 for each genotype), water volume (J, n=40 for each genotype) and of volumes of extractable hemolymph (K, n=40 for each genotype) from larvae reared on standard food versus food made from *erg2Δ* yeast. The dotted line represents the mean. **p<0,01, ***p<0,001. (L) Magnified larvae reared in food containing the erg2 mutant yeast strain *erg2Δ*. All larvae-bearing tumors showed swelling defects independently of their genetic background. (M,N) Tumor sizes (M, n=30 per genotype) and aggressiveness (N, n=40 for each genotype) in larvae with the indicated genotypes reared on standard food versus food containing *erg2Δ* yeast. The dotted line represents the mean. *p<0,05, **p<0,01. (O) Differences in dry mass observed in larvae with the indicated genotypes fed with standard or *erg2Δ*-food (n=40 for each genotype). The delta between *CTRL>* and *tumor>* is plotted as a single value. ***p<0,001.

**Figure 4.**
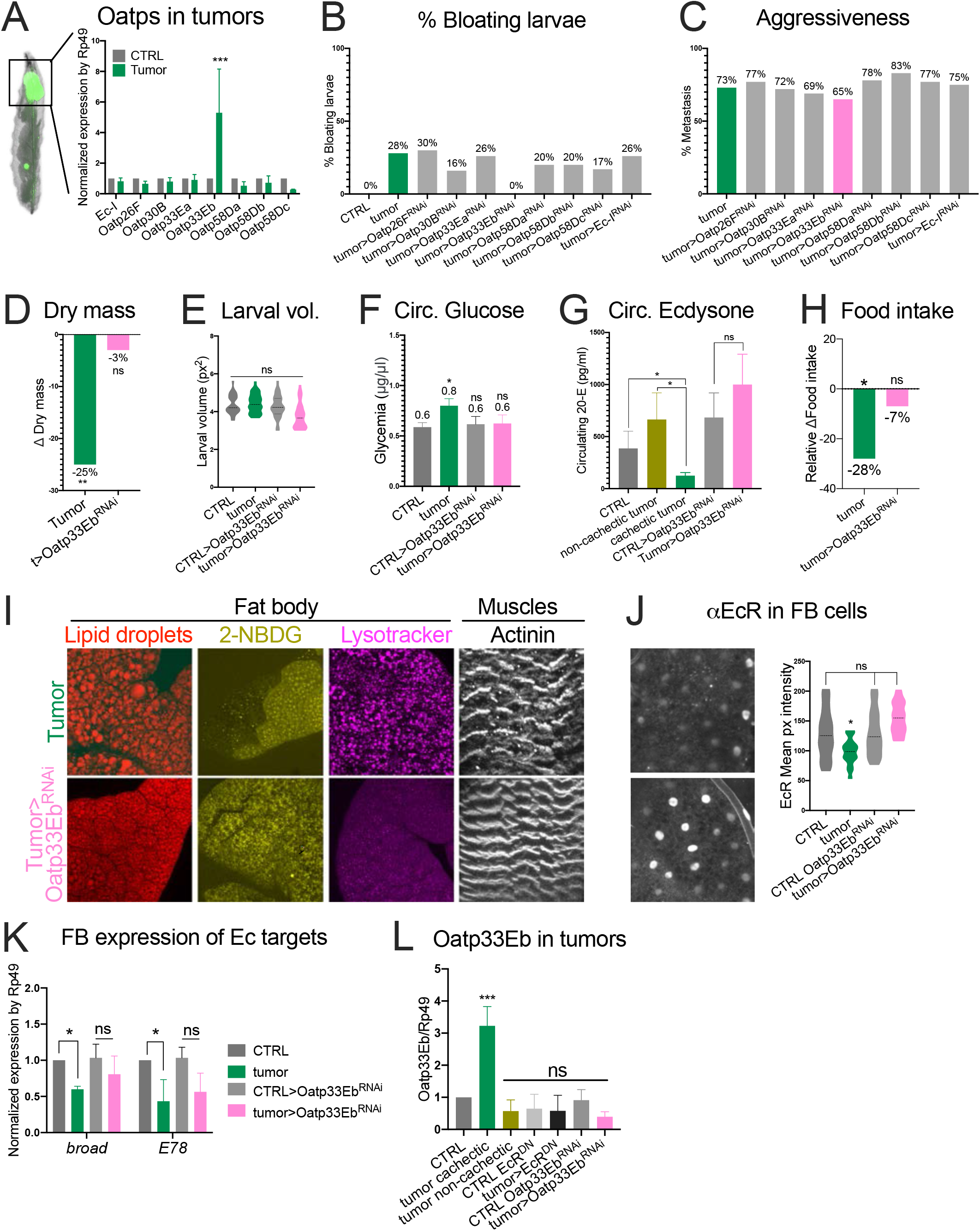
The *SLCO* transporter *Oapt33Eb* is upregulated in the tumor and drives the sink effect. (A) mRNA levels of *Drosophila Oatp* genes from CTRL and tumorous discs measured by RT-PCR. Shown results are mean +/-SEM. ***p<0,001. (B, C) Percentage of larvae with bloating defects after the individual inhibition of each *Oatp* gene with an RNAi line (B, n=200 for each genotype) and metastasis (C, n=200 for each genotype). (D-G) Differences in dry mass (D, n=50 for each genotype), larval volume (E, n=50 for each genotype), circulating glucose (F, mean +/-SEM) and circulating levels of 20-E (G) in larvae with the indicated genotype. *p<0,05, **p<0,01. (H) Food intake of larvae with the indicated genotypes (n=11 groups of 8 larvae each one). Values are shown relative to control, *p<0,05. (I) Fat bodies and muscles from tumors and *tumors>Oatp33Eb*^*RNAi*^ animals. Lipid droplets are shown in red, Glucose uptake in yellow, acidic vesicles in purple and muscle fibers in white. (J) EcR staining in fat body cells from tumors and *tumors>Oatp33Eb*^*RNAi*^ animals. EcR is significantly reduced in *tumors>* but not *tumors>Oatp33Eb*^*RNAi*^ fat bodies. The dotted line represents the mean. *p<0,05. (K) mRNA levels of ecdysone targets (*broad* and *E78*) measured by RT-PCR in the fat body of animals with the indicated genotypes. The results shown are mean +/-SEM. *p<0,05. *(L) Oatp33Eb* expression in different tumors measured by RT-PCR (mean +/-SEM). ***p<0,001.

### The transporter *Oapt33Eb* is upregulated in the tumor and drives the sink effect

Since *Ec-I/Oatp74D* drives ecdysone uptake in the tumors when over-expressed, we hypothesized that its expression could be upregulated in tumorous discs compared to normal discs, but this is not the case (Fig. 4A). Moreover, silencing *Ec-I* in the tumor did not rescue cachectic syndromes (Fig. 4B, Suppl. Fig 5A), showing that *Ec-I* is not responsible for the sink effect. Seven other Oatp-encoding genes are present in the fly genome and we compared their expression in normal and tumorous discs. Only *Oatp33Eb* was found upregulated in tumor tissues (Fig. 4A). When targeted to tumor cells, only *Oatp33Eb-RNAi* could prevent cachexia and restore normal levels of circulating ecdysone without modifying the tumor burden or aggressiveness (Fig. 4B-I, Suppl. Fig 5B-G). Accordingly, in fat cells of *tumor>Oatp33Eb*^*RNAi animals*^, the levels of EcR target genes as well as EcR antibody staining were rescued (Fig. 4J,K). These results indicate that *Oatp33Eb* is the only gene of the family to be upregulated in the tumor and that this upregulation is responsible for the observed sink effect and the subsequent cachectic syndrome. Notably, expression of *shade* is not upregulated in tumor cells, indicating that they do not modulate their ability to convert E into 20E (Suppl. Fig 5H).

**Figure 5.**
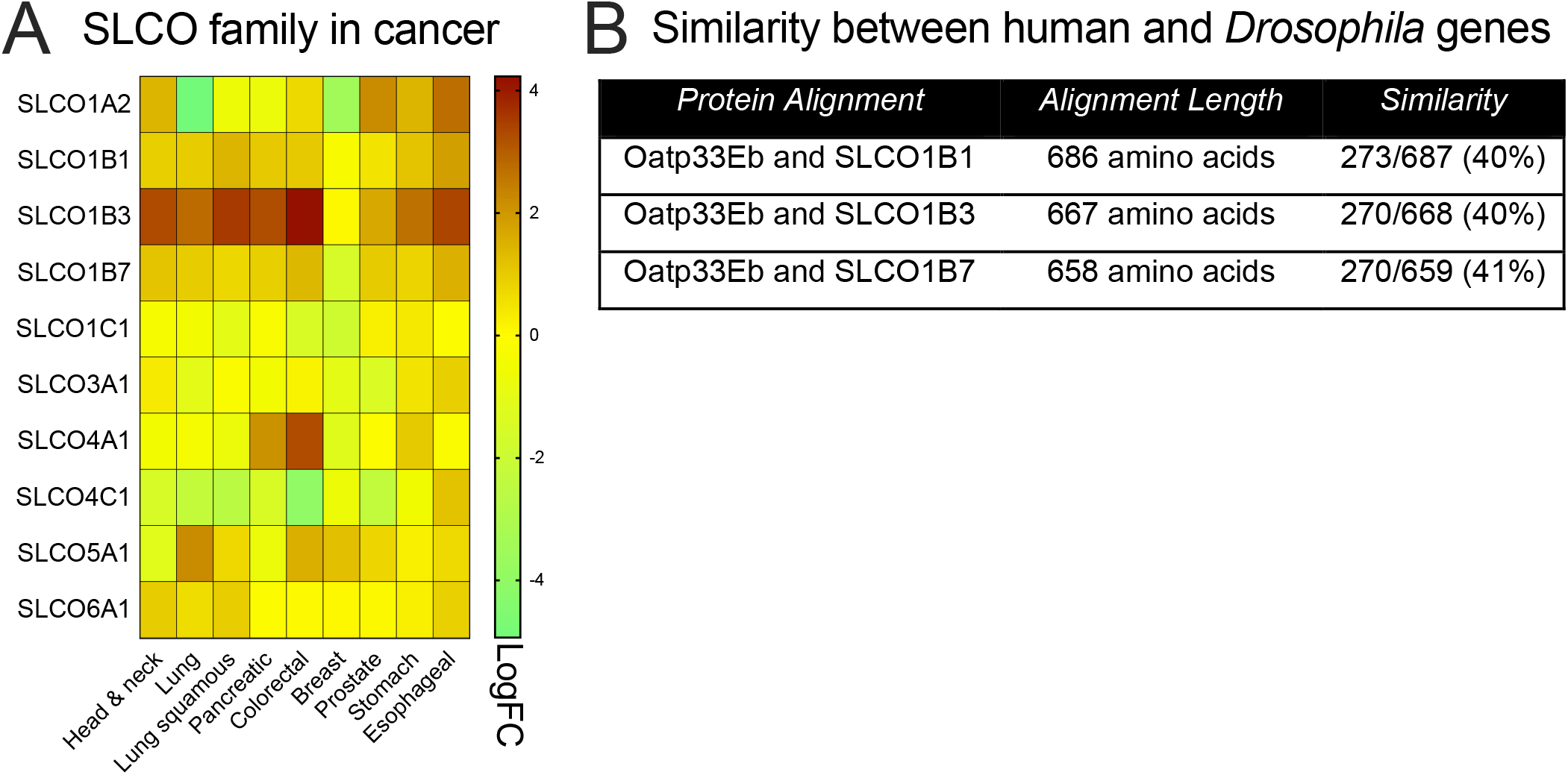
SLCO family genes upregulated in human cancer. (A) Heatmap showing the *SLCO* family genes differentially expressed in normal *vs* tumorous tissues. Intensities were Log2 transformed and are displayed in colors ranging from burgundy to light green as shown in the key. Each row represents a gene and each column a type of cancer. (B) Amino-acid sequence comparison between *Drosophila Oatp33Eb* and human *SLCO1B1, SLCO1B3* and *SLCO1B7* genes. (C) An integrative model of the presented work. We propose that the Oapt33Eb transporter triggers the sink effect in the tumor. This titers circulating 20-E available for peripheral tissues. The lack of ecdysone generates a catabolic state, eventually promoting cachexia. Molecules produced by the tumor called ‘cachectokines’ could also influence this process.

Oatp33Eb appears to act specifically on the levels of circulating ecdysone. Indeed, ecdysteroidogenic gene expression was not found altered in *tumor>Oatp33Eb*^*RNAi*^, neither PTTH nor Serotonin circuitries (Suppl. Fig 5I,J). *tumor>Oatp33Eb*^*RNAi*^ larvae where slightly advanced compared to *tumor>* larvae and still produced Dilp8 (Suppl. Fig 5L, M).

Finally, we found that expression of *Oatp33Eb* is subjected to strong feedbacks by EcR signaling. Indeed, inhibition of EcR function (*tumor>EcR*^*DN*^) prevented the *Oatp33Eb* upregulation observed in tumor cells (Fig. 4L). This is in line with the high number of putative EcR/usp binding sites present in the promoter region of the Oatp33Eb gene (Suppl. Fig5N). This result could explain the suppression of cachexia observed upon inhibition of intra-tumor EcR signaling (see Fig. 1, Suppl. Fig. 1 and 2).

Oatp channels are conserved in human (*solute carrier organic anion* -*SLCO-*gene family) and were previously shown to transport xenobiotics and drugs such as anticancer agents, and possibly hormones (Obaidat, Roth and Hagenbuch, 2012). Searching the *TGCA* database, we found that some of the *SLCO* genes are preferentially expressed in tumors with high prevalence of cachexia (advanced head and neck, 57% prevalence; lung non-small cell, 36% prevalence; pancreatic, 40-54% prevalence; colorectal, 28% prevalence; breast, 30% prevalence; prostate; 50% prevalence) (Fig. 5A). *SLCO1B3, SLCO1B1* and *SLCO1B7*, the most upregulated genes of the family, show highest similarity (40%) with *Drosophila Oatp33Eb* (Fig. 5B). This and our findings in the fly model suggest a possible conservation of the role of human OATPs in steroid depletion in cancer-induced cachexia.

## Discussion

The *Drosophila* larva has recently emerged as a genetic model to study several aspects of normal physiology. Here we use this model to study pathophysiological conditions linked to tumor development. The prevalence of cachexia is dramatically increased in the larval population when Ec-I, one member of *Drosophila* Oatp transporters, is upregulated in tumors. However, upregulating Ec-I in non-tumorous discs is not sufficient to promote a catabolic conversion. This suggests that additional pro-cachectic, tumor-derived factors are required to promote the cachectic state, as recently established using the adult *Drosophila* model. Among the previously described pro-cachectic factors, Impl2 is found up-regulated in cachectic tumors. However, knocking down Impl-L2 in the tumors does not reduce the prevalence of cachexia in larvae, suggesting that it does not play a major role in triggering cachexia in our model. Neither Pvf1, nor branchless (bnl) are upregulated in tumors in our conditions (see Suppl. Fig. 2G).

Our finding that cachectic tumor-bearing animals have reduced levels of the steroid ecdysone parallels the prevalence of hypogonadism in men with cancer cachexia (Burney *et al*., 2012). Remarkably, the capacity of cachectic animals to produce ecdysone or to convert it into the active form 20-E seems unaffected and this reduction mostly results from an increase in steroid uptake by the tumor.

Interestingly, the gene encoding Oatp74D, recently described as Ecdysone Importer (Ec-I), is not induced in cachectic tumors and its silencing does not reduce the prevalence of cachexia. Rather, we find that another member of the Oatp family, Oatp33Eb, is specifically induced in cachectic tumors. The link between this Oatp member and ecdysone import is not molecularly established. However, we find that silencing Oatp33Eb efficiently rescues both cachexia and the levels of circulating ecdysone. Moreover, silencing EcR in tumors prevents *Oatp33Eb* upregulation (see Fig. 4L), suggesting that EcR signaling exerts a positive feedback on its expression, in line with numerous EcR/Usp binding sites found in the *oatp33Eb* promoter. This could explain the efficient rescue of cachexia observed when EcR signaling is inhibited in tumors (*tumor>EcR*^*DN*^). Indeed, in these conditions, not only Oatp33Eb is down regulated, but the circulating levels of ecdysone are rescued to normal (our Fig. 1K). Interestingly, growing these animals on *erg2?* food was sufficient to trigger cachexia in all larvae, indicating that reducing circulating levels of ecdysone is the limiting trigger for cachectic conversion. A similar feedback mechanism has been described in the case of the vertebrate *Oatp1* gene, whose expression is controlled by the level of androgen in the kidney (Isern *et al*., 2001).

The role of human OATPs in transporting xenobiotics has made them possible markers for chemotherapy disposition (Schulte and Ho, 2019). Several evidences in cell and tumor models also indicate a possible role for OATP1B3 (the closest vertebrate homolog of the fly Oatp33Eb) in transporting steroid hormones (Hamada *et al*., 2008). Some of these transporters are highly expressed in tumors associated with high prevalence of cachexia including breast, colon, prostate, pancreatic and head & neck cancers, with little expression in normal tissues. This mimics the situation observed with Oatp33Eb in our tumor model and suggests that OATPs could serve as biomarkers or therapeutic targets for the early stages of cancer-induced cachexia.

## Acknowledgements

We thank Mike O’Connor, and lab members for insightful discussions and comments on the manuscript. We thank the Vienna Drosophila RNAi Center and the Bloomington stock center for lines. We thank the PICT-IBiSA@BDD light-microscopy facility of Institut Curie. This work was supported by Institut Curie, CNRS, INSERM, ARC Fondation (grant n° PDF20180507272 to P.S.), FRM, European Research Council (Advanced Grant n°694677 to P.L.), and the Labex DEEP program (ANR-11-LABX-0044, ANR-10-IDEX-0001-02).

## Author contributions

Conceptualization, P.L. and P.S., Methodology, P.L. and P.S., Investigation, P.S., Writing and editing, P.L. and P.S., Funding Acquisition, P.L. and P.S.; Supervision, P.L.

## STAR METHODS

### CONTACT FOR REAGENT AND RESOURCE SHARING

Further information and requests for resources and reagents should be directed to and will be fulfilled by the Lead Contact, Pierre Léopold.

### EXPERIMENTAL MODEL AND SUBJECT DETAILS

#### Drosophila strains and maintenance

Animals were reared at 25°C on fly food containing, per liter: 50g yeast powder, 35g wheat flour, 7.5 g agar, 55g sugar, 25ml Methyl and 4ml propionic acid.

*Drosophila* strains used in this work were *w*^*118*^, *eyFLP, act>FRT-y*^*+*^*-FRT>Gal4, UAS-GFP, UAS-Ras*^*v12*^, *UAS-dlg*^*RNAi*^ from BDSC. *UAS-Ec-I* (Okamoto *et al*., 2018b), *UAS-EcR*^*DN*^ (Cherbas *et al*., 2003), *UAS-shd*^*RNAi*^ (BDSC 67356), *UAS-Cyp18a1* (Rewitz, Yamanaka and O’Connor, 2010a) *tGPH* (Britton *et al*., 2002), *EcRE_GFP* (Kimura *et al*., 2008), *UAS-Ec-I*^*RNAi*^ (VDRC 37295), *UAS-Oatp26F*^*RNAi*^ (VDRC 2650), *UAS-Oatp30B*^*RNAi*^ (VDRC 22983), *UAS-Oatp33Ea*^*RNAi*^ (VDRC 45896), *UAS-Oatp33Eb*^*RNAi*^ (VDRC 100431), *UAS-Oatp58Da*^*RNAi*^ (VDRC 6623), *UAS-Oatp58Db*^*RNAi*^ (VDRC 100348) and *UAS-Oatp58Dc*^*RNAi*^ (VDRC 39469).

Flies were reared and experiments were performed on fly food containing, per liter: 50g yeast powder, 35g wheat flour, 7.5 g agar, 55g sugar, 25ml Methyl and 4ml propionic acid. Experiments were done at 25°C. For all experiments, both males and females were used. For all experiments, a precise staging of the animals was done with 4-hours egg layings collected on agar plates with yeast. The next day, recently hatched L1 larvae were collected 24 hours AEL and reared in tubes (forty larvae each) containing standard food. The developmental stage or time of development at which analysis were done is indicated in the methods section.

## METHODS DETAILS

### Tumor generation

We induced the formation of metastatic tumors as described in Pagliarini and Xu, 2003. We utilized *eyeless* promoter–driven FLP recombinase expression (*eyFLP*) to activate the *act>y+>Gal4* construct. This allowed the expression of the oncogenic *ras*^*V12*^ mutant with an interference RNA to inhibit the tumor suppressor gene *disc large* (*dlg*^*RNAi*^) into GFP-labeled cells specifically in the developing larval eye-antennal imaginal discs, which altogether is known to induce cachexia in *Drosophila* adults after transplantation (Figueroa-Clarevega and Bilder, 2015; Kwon *et al*., 2015). Females containing *eyFLP; UAS-Ras*^*v12*^, *UAS-dlg*^*RNAi*^*/CyO*_*Gal80*_^*TS*^; *act>y*^*+*^*>Gal4, UAS-GFP/TM6B* were crossed with the desired males. For each genotype, we used as a control larvae from the same vial but tumor-free (*eyFLP; +/CyO*_*Gal80*_^*TS*^; *act>y*^*+*^*>Gal4, UAS-GFP/+)*. Tumor size was analyzed at 120h AEL with Fiji software. The number of metastasis was counted under a GFP scope.

### Modification of ecdysone signaling

We used distinct tools to modify ecdysone signaling in peripheral tissues. EcR^DN^ is a dominant negative form of EcR, which binds to the promoter region of target genes but is deficient for transcriptional activation (Cherbas *et al*., 2003). The *shade* (*shd*)*-RNAi* construct blocks expression of *shade*, a gene coding for a P450 enzyme that converts ecdysone in its active form 20-hydroxyecdysone (Petryk *et al*., 2003). The *Cyp18a1* gene codes for an enzyme that degrades ecdysone (Rewitz, Yamanaka and O’Connor, 2010a). Ec-I is an Oatp importer (Oatp74D) that transports ecdysone into *Drosophila* cells (Okamoto *et al*., 2018b).

### Test for bloating larvae and statistics

Late third instar larvae (120h AEL) were put in PBS and imaged under a microscope. We considered a ‘bloating larvae’ when they were inflated and we detected transparent regions that were usually filled with fat body content. Therefore, the % of bloating larvae was calculated after the number of larvae that showed this phenotype (n=200 for each genotype). For each sample we scored the percentage of individuals that belong to the **“**bloating larvae**”** category and calculated the standard error of sample proportion based on binomial distribution (bloating or not) SE = √ p (1-p)/n, where p is the proportion of successes in the population. Volume was calculated using Fiji software.

### Water volume, hemolymph volume and dry mass

We adapted the protocol to *Drosophila* larvae from the one described in Folk, Han and Bradley, 2001. Synchronized third instar larvae were first rinsed in PBS 1X and dried in a Kimwipe paper. Afterwards they were weighted in groups of 10 (total mass). Hemolymph was blotted out from the abdominal opening with a Kimwipe moistened with PBS1X. Each group of larvae were reweighted (total mass-hemolymph content), dried for 1 hour at 60° C and weighted a third time (dry mass). The water accumulation was computed by subtracting the dry mass from the wet mass. The hemolymph content was estimated by determining the reduction in mass following hemolymph blotting. The percentage of dry mass was calculated with the formula dry mass*100/wet mass. Delta Δ dry mass shows variation in total dry mass between control and tumor-bearing larvae. The delta between the two conditions is plotted as a single value. For each experiment, 5 replicates were carried out. P values were calculated after a one-way ANOVA with the Tukey-Kramer multiple comparisons test.

### Food intake

Food intake was calculated as described in Bjordal *et al*., 2014. Synchronized pre-wandering third instar larvae were washed in PBS 1X, dried on a Kimwipe and incubated for 3 hours at 25° C in a petri dish with regular food supplemented with 1,5% w/v of blue dye (Erioglaucine Disodium Salt, Sigma-Aldrich). Afterwards, larvae were washed again, dried and put together in groups of 8 in eppendorf tubes to be frozen in liquid N_2_. Then they were transferred to a freezer at -20° C. Samples were homogenized in 20ul PBS 1X and centrifuged for 20 min at 4° C. 10ul were finally put in a new tube and the amount of blue dye in the supernatant was measured spectrophotometrically (OD_629nm_). Relative Δ food intake shows variations in food intake according to different tumor-bearing larvae compared with their controls, respectively. The delta between the two conditions is plotted as a single value. P values were calculated after an ordinary one-way ANOVA test. The quantification of the mean larval weight was carried out immediately after this protocol and P values were calculated after a one-way ANOVA with the Tukey-Kramer multiple comparisons test.

### Immunochemistry

Immunostaining and was performed using standard protocols. Primary antibodies used in this work were αGFP (1:10.000 Abcam), αactinin (1:100 DHSB 2G3-3D7) Phalloidin-FITC (1:200 Sigma P-5282), αEcR (1:30 DHSB Ag10.2), αSerotonin (1:500 Sigma-Aldrich S5545), αPtth (1:200, our lab). Fluorescently labeled secondary antibodies were from Thermofisher Scientific. Larval tissues were mounted in Slowfade™ Diamond with DAPI (Thermofisher) and images were taken under an inverted Laser Scanning Confocal Microscope with Spectral Detection (LSM900 - Zeiss).

#### Lysotracker Staining

We followed the protocol described in Devorkin and Gorski, 2014. Fat bodies from third instar larvae were dissected in PBS and then incubated for 5 minutes in 100 μM LysoTracker Red DND-99 (Invitrogen) at RT. Then they were fixed in 4% paraformaldehyde for 30 minutes, washed in PBS and mounted in Slowfade™ Diamond (Thermofisher).

#### Nile Red Staining

Fat bodies from third instar larvae were dissected in PBS and immediately fixed in 8% paraformaldehyde for 45 minutes. They were washed twice in PBTween 0,1% and incubated for 5 minutes in diluted Nile Red Solution (1:8000 from 200ug/ml stock in DMSO, Sigma N3013) at RT. They were washed again in PBTween 0,1% and mounted in Slowfade™ Diamond (Thermofisher). Fiji software was used to quantify lipid droplets and droplet size.

#### 2-NBDG glucose analog staining

The dye was reconstituted with 200μl RNAase-free water and kept at -20° C. Then fat bodies were incubated in 2-NBDG diluted 1:100 in PBX1X for 15 minutes in the dark, rinsed twice and fixed in 4% paraformaldehyde for 30 minutes at RT. Tissues were rinsed again and mounted in Prolong™ (Thermofisher).

### Quantitative RT-PCR

Larvae were collected at the indicated time point AEL and specific tissues were dissected in cold PBS1X then frozen immediately in liquid N_2_. Total RNA was extracted using a QIAGEN RNeasy lipid tissue minikit or microkit according to the manufacturer’s protocol. 2μg of each sample were treated with DNAse I (invitrogen) and reversely transcribed using a SuperScript II reverse transcriptase (Invitrogen). The resulting cDNA was utilized for quantitative qPCR (Viia 7; Applied Biosystems). cDNA was diluted 1:50 in H_2_0 and mixed with 10uM primers and Power SYBR™ Green PCR Master Mix (Applied Biosystems). Samples were normalized to levels of *ribosomal protein (rp)49* transcript levels. Three separate biological samples were collected for each experiment and triplicate measurements were conducted.

Primers used in this work:

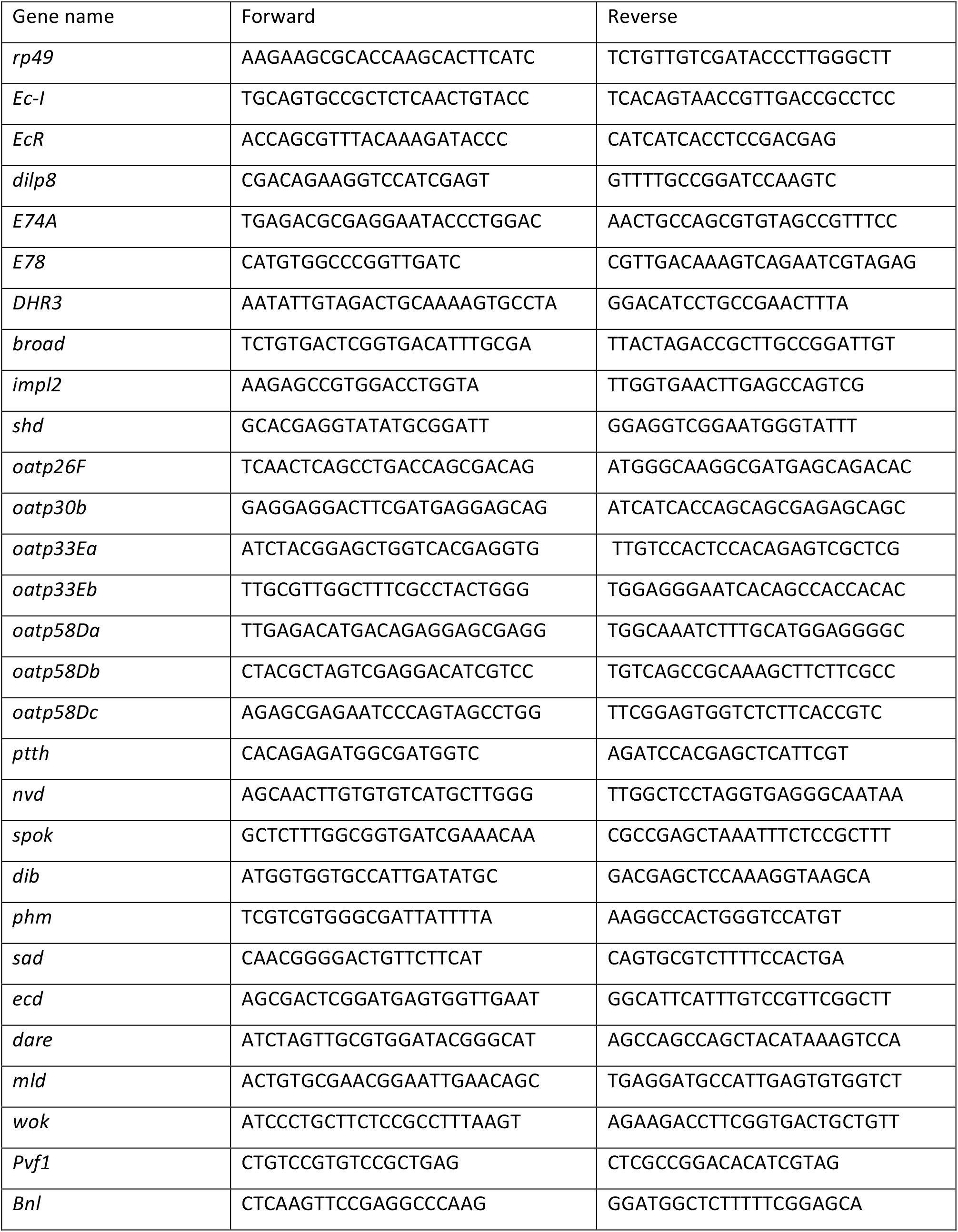

### Pupariation curves

4h egg-laids were carried out in small plates made of PBS 1X, 2% agar and 2% Glucose. After 24 hours first instar larvae were transferred to vials filled with standard fly food (40-50 larvae per tube) and let in an incubator at 25° C. The number of larvae that had pupariated at a given time AEL was scored at 114, 120, 128, 132, 140, 144, 152, 156 and 164 hours.

### Circulating ecdysone measurements

5ul of hemolymph for each genotype were homogenized and extracted in 200ul methanol. Extractions were then submitted to a competitive ELISA test as previously published (Rewitz, Yamanaka and O’Connor, 2010b) to evaluate the amount of circulating 20-E, using the detection kit from SpiBio (Bertin reagents) as recommended by the manufacturer. Absorbance at 415nm was detected using a TECAN microplate reader.

### Trehalose and Glucose measurements

For each genotype, synchronized first instar larvae larvae were incubated at 25**°** C until early third instar. Three technical replicates of 1,5ul hemolymph were used for each condition. Hemolymph samples were diluted in 149ul test buffer TB (5mM Tris pH6.6, 137mM NaCl and 2.7mM KCl) and heated for 5 min at 70**°** C. Each sample was separated into two tubes of 70ul. 70ul of TB were added to one of the tubes and 70ul of trehalose buffer TbT (1mL TB + 3uL Porcine trehalase, Sigma, T8778-1UN) to the other. Trehalose and Glucose standards (1, 0,16, 0,08, 0,04, 0,02 and 0,01 mg/ml, Trehalose from Sigma 90208) were prepared in the same way. Both the samples and the standards were incubated at 37° C overnight. The next day tubes were centrifuged for 3 minutes and 30ul of each one was transferred to individual wells of a 96-well plate. 100ul of GO reagent (GAGO-20 Sigma) were put on and then the plate was sealed and incubated at 37° C for 30 to 60 minutes. To stop the reaction, 100ul of sulfuric acid acid were added. Absorbance at 540nm was detected using a TECAN microplate reader. Concentration was calculated based on the equation given by a linear regression and the dilution. P values were calculated after an ordinary one-way ANOVA test.

### *Ex-vivo* incubation of fat bodies

Fat bodies from *tGPH* larvae with indicated genotypes were incubated for 30 minutes at RT in Schneider’s medium with or without human insulin (0,5uM #I9278 Sigma). Incubation mix was then removed and tissues were fixed and processed as described above. Chicken anti-GFP antibody (1:10000, Abcam) was used as primary. Images were obtained under an inverted Laser Scanning Confocal Microscope with Spectral Detection (LSM900 - Zeiss) and processed with Fiji Software. Delta tGPH shows variation in membrane tGPH between fat bodies from the same larvae incubated with or without insulin. The delta between the two conditions is plotted as a single value. P values were calculated after a one-way ANOVA with the Tukey-Kramer multiple comparisons test.

### Preparation of *erg2Δ::Trp1* yeast food

Fly food made of *erg2Δ::Trp1* yeast was prepared as previously described (Katsuyama and Paro, 2013).

#### Yeast culture

A synthetic complete tryptophan-minus medium was prepared and autoclaved (6.7g yeast nitrogen base without amino acids BD DIFCO 291940, 5g Cassamino acids BD Bacto 223050, 20g Glucose Sigma G7021 and 1L miliQ water). 10ml of 100X Uracil were added and the *erg2Δ::Trp1* yeast was inoculated and cultured at 37° C until reaching an OD_600nm_=1. Then the harvested cells were centrifuged and washed several times with PBS and the clean 10ml yeast pellet was immediately used or stored frozen until used.

#### erg2Δ food preparation

The frozen 10ml yeast pellet was suspended in 10ml PBS 1X and boiled for 30 minutes. Then the heat-inactivated yeast paste was mixed with 0.1g glucose, 0.12 agar, 10 ml water and 300ul of propionic acid under sterilized conditions. The food was distributed in vials (5ml per vial).

#### erg2Δ food experimental conditions

Staged embryos were sterilized with 3% sodium hypochlorite solution, washed twice in PBS 1X and transferred to vials with the help of a sterilized net (around 25 eggs per vial). The tubes were transferred to a 25 degrees chamber and larvae were analyzed at specific time points (120h AEL).

### 20-E treatment

4h egg layings were carried out in small plates made of PBS 1X, 2% agar and 2% Glucose. After 24 hours first instar larvae were transferred to vials filled with standard fly food (40-50 larvae per tube) and let in an incubator at 25° C. At 48, 72 and 96h AEL, 100ul of a 0,2mg /ml ethanol of 20-E (Sigma, H5142) fresh solution were added, and larvae were analyzed at 120h AEL. In parallel, we added the same quantity of ethanol to control tubes.

### Binding site prediction

*Oatp33Eb* genomic sequence was taken from the NCBI reference sequences collection (RefSeq). Putative Binding sites were analyzed in JASPAR database (http://jaspar2016.genereg.net/) using the EcR::Usp matrix model (MA0534.1, Uniprot ID: P34021 P20153, PubMed ID 8649409) with a Relative profile score threshold of 80%.

