## Supplementary information for "An Oatp transporter-mediated steroid sink promotes tumor-induced cachexia in *Drosophila*"

**SUPPLEMENTAL INFORMATION**

### Supplemental Figure 1

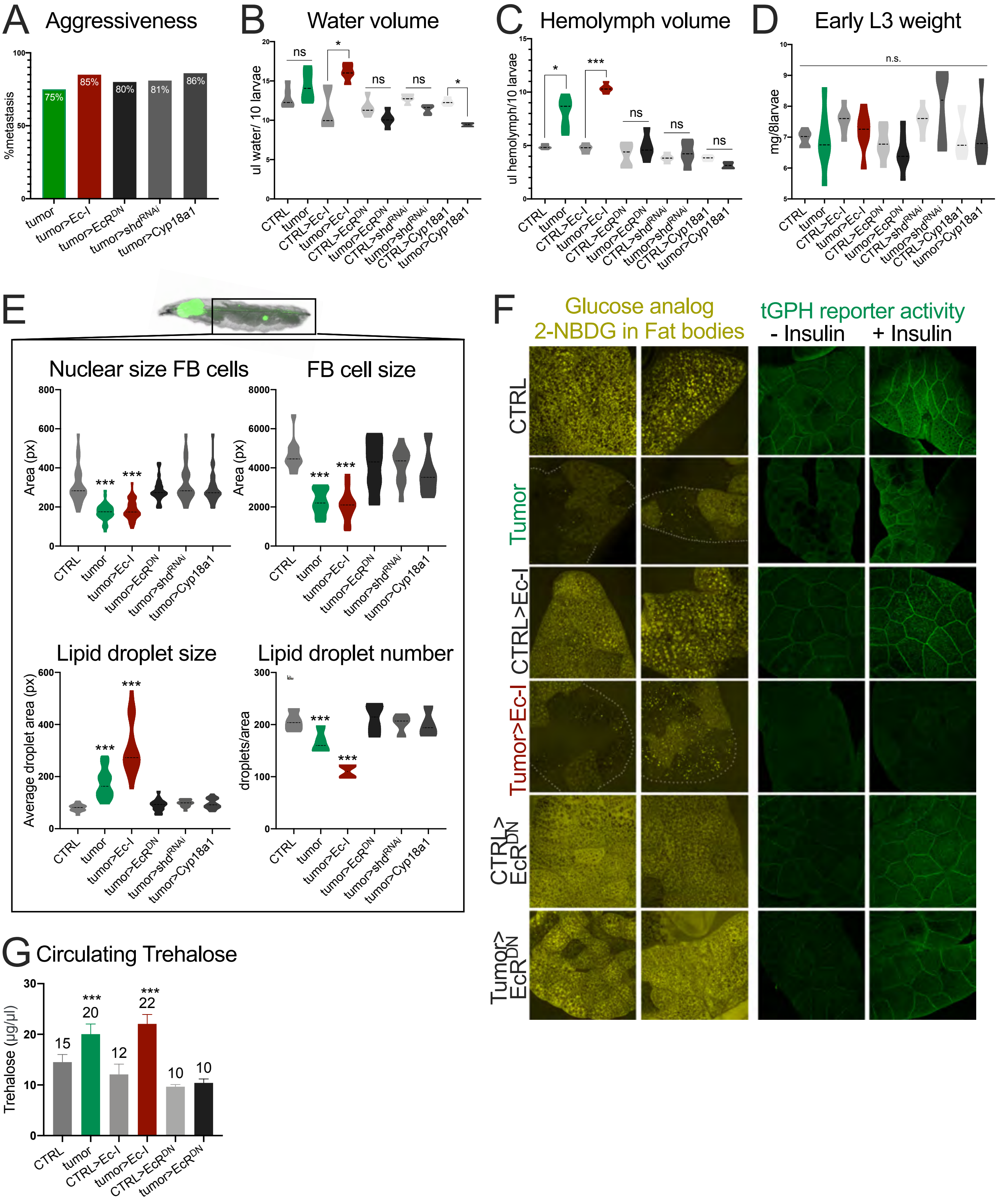

**Supplemental Figure 1. Supporting data for Fig. 1 (I).**

(A) Percentage of larvae with the indicated genotype with metastasis (n=200 for each genotype).

(B, C) Water volume and the volumes of extractable hemolymph measured in larvae (n=50 for each genotype). The dotted line represents the mean. *p<0,05, ***p<0,001.

(D) Larval weight just after the food intake assay (n=12 groups of 8 larvae each one). No significant differences were observed.

(E) Quantification of different parameters measured in fat body cells from larvae with the indicated genotypes: Size of the nuclei (n=50), cell size (n=50), lipid droplet size (n=50) and lipid droplet number (5 groups in different areas). *p<0,05, ***p<0,001. The dotted line represents the mean.

(F) Left: fat bodies dissected from the indicated genotypes and incubated with the glucose analog 2-NBDG (in yellow). Right: fat bodies showing the *tGPH* reporter with or without insulin incubation (in green). The quantification is shown in Fig. 1H.

(G) Concentration of Trehalose in the hemolymph from larvae with the indicated genotypes (mean +/- SEM, ***p<0,001).

### Supplemental Figure 2

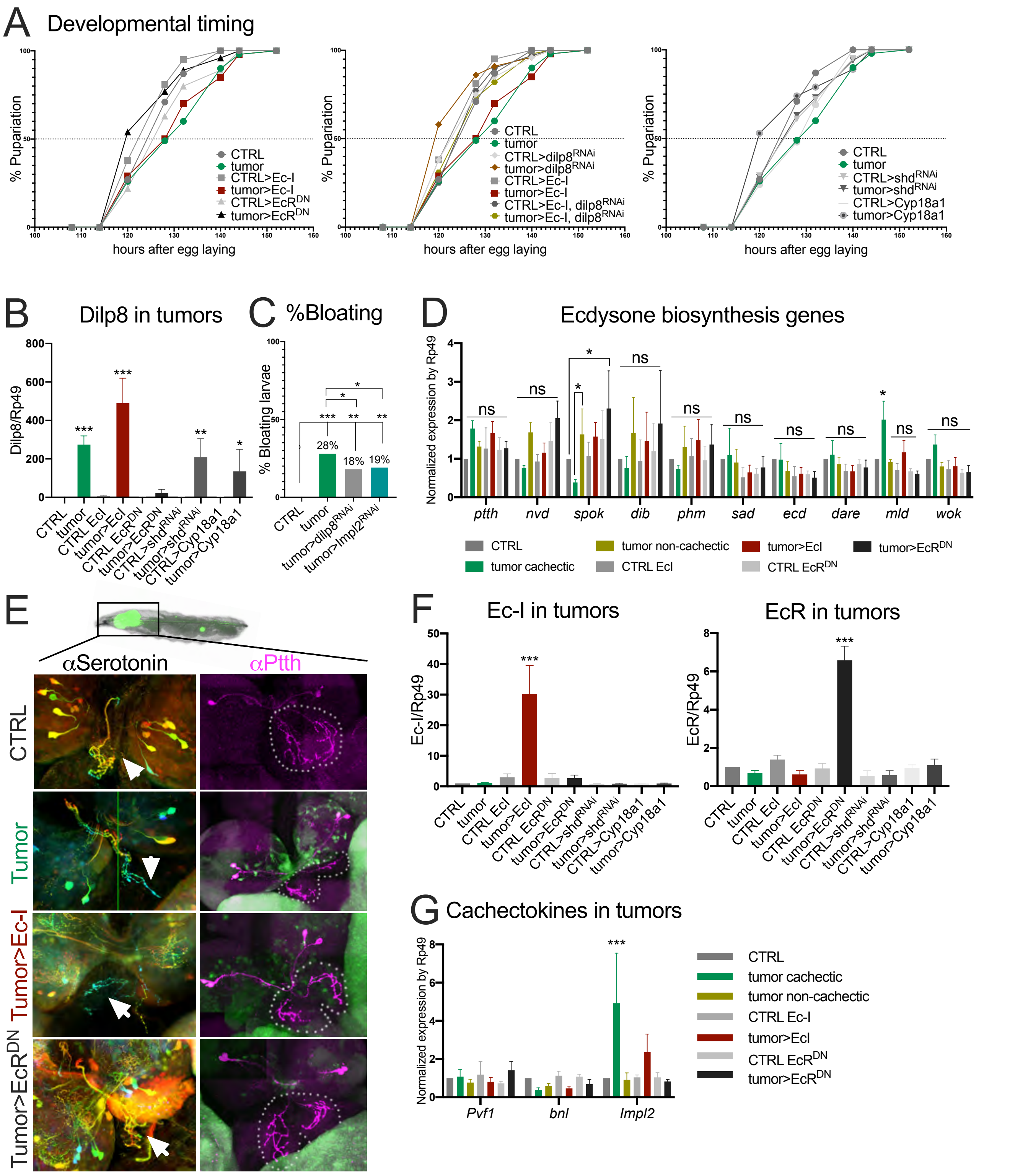

**Supplemental Figure 2. Supporting data for Fig. 1 (II).**

(A) Pupariation timing after tumor induction. Larvae-bearing tumors and *tumors>Ec-I* showed a 6-8 hours delay that is rescued by *dilp8* inhibition. Larvae bearing *tumors>EcR$^{DN}$*, *>shd$^{RNAi}$* and *>Cyp18a1* did not show pupariation delay.

(B) *Dilp8* expression in CTRL or tumor> discs with the indicated genotypes measured by RT-PCR. *p<0,05, **p<0,05, ***p<0,001.

(C) Larvae still show bloating defects after *dilp8-* and *impl2*-RNAi silencing. *p<0,05, **p<0,05, ***p<0,001.

(D) mRNA levels of the genes involved in ecdysone synthesis in different types of tumors measured by RT-PCR. The results shown are mean +/- SEM. *mld* is upregulated in cachectic tumors. However, this upregulation persists in *tumor>Oatp33Eb-RNAi* animals, where cachexia is fully rescued (see Suppl. Fig. 5J). *p<0,05, **p<0,01, ***p<0,001.

(E) Serotonin and Ptth stainings in brains and prothoracic glands (PGs) of larvae-bearing tumors. The white arrows point to neuronal projections into the PG and the white dotted line surrounds the PG. No morphological differences were seen among different genotypes.

(F,G) mRNA levels of *Ec-I*, *EcR* and the cachectic inducers *Pvf1*, *bnl* and *Impl2* from different types of tumors measured by RT-PCR. The results shown are mean +/- SEM. ***p<0,001.

### Supplemental Figure 3

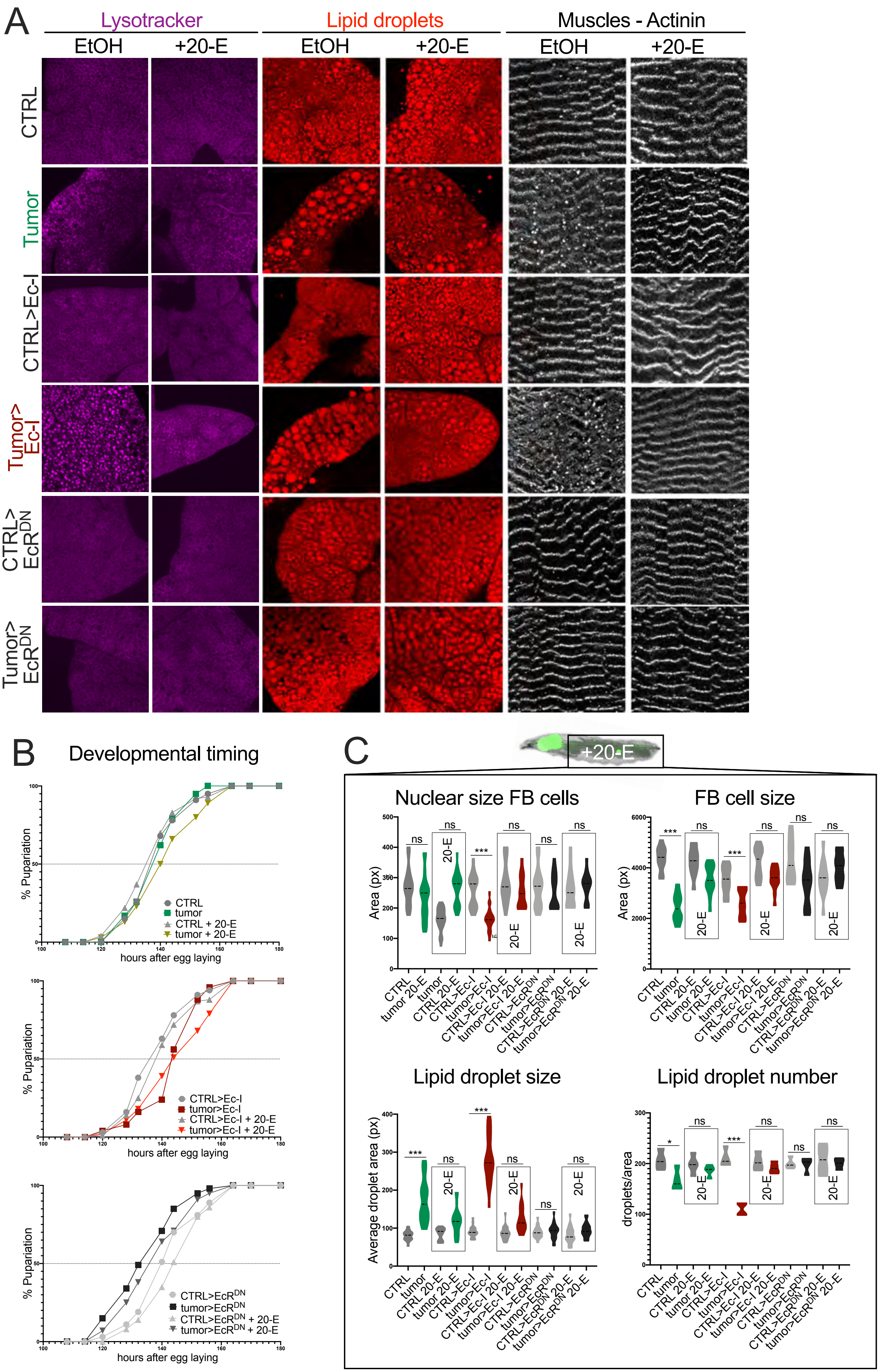

**Supplemental Figure 3. Supporting information for Figure 3.**

(A) Peripheral tissue wasting in larvae reared on standard (EtOH: ethanol) and 20-E-food (20-E diluted in ethanol). Acidic vesicles are labeled with Lysotracker (purple), lipid droplets with Nile Red (red) and muscle fibers with α-actinin (white). The addition of 20-E into the food improves the cachectic phenotype both in fat bodies and muscles.

(B) Effects on the developmental timing after tumor formation at the indicated time points in larvae reared on standard food versus food containing exogenous 20-E. Pupariation is not advanced in any case.

(C) Parameters measured in fat body cells from larvae with the indicated genotypes: Size of the nuclei (n=50), cell size (n=50), lipid droplet size (n=50) and lipid droplet number (5 groups in different areas). Larvae fed with food containing exogenous 20-E are compared with larvae reared on standard food in parallel. Grey squares highlight data from larvae treated with 20-E. The dotted lines represent the mean. ***p<0,001.

### Supplemental Figure 4

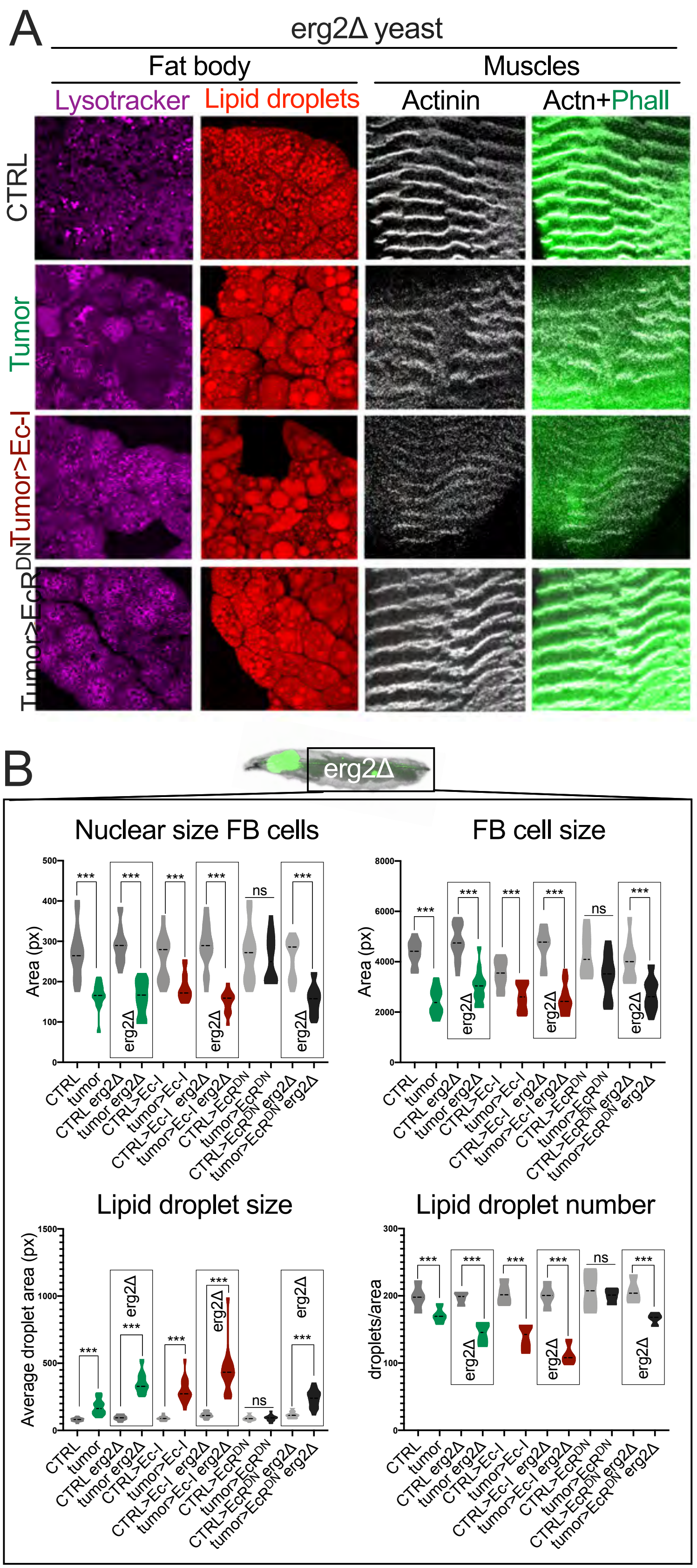

**Supplemental Figure 4. Supporting information for Fig. 3.**

(A) Peripheral tissue wasting is aggravated in in larvae reared on food containing *erg2Δ* yeast. Acidic vesicles are labeled with Lysotracker (purple), lipid droplets with Nile Red (red) and muscle fibers with α-actinin (white) and phalloidin (green). When fed with *erg2Δ* yeast, all larvae-bearing tumors showed an increase in autophagy and larger lipid droplets, and all except *tumor>EcR*$^{DN}$ displayed signs of muscle fiber degradation.

(B) Parameters measured in fat body cells from larvae with the indicated genotypes: Size of the nuclei (n=50), cell size (n=50), lipid droplet size (n=50) and lipid droplet number (5 groups in different areas). The dotted line represents the mean. Larvae fed with *erg2Δ* yeast are compared with larvae reared on standard food in parallel. ***p<0,001.

### Supplemental Figure 5

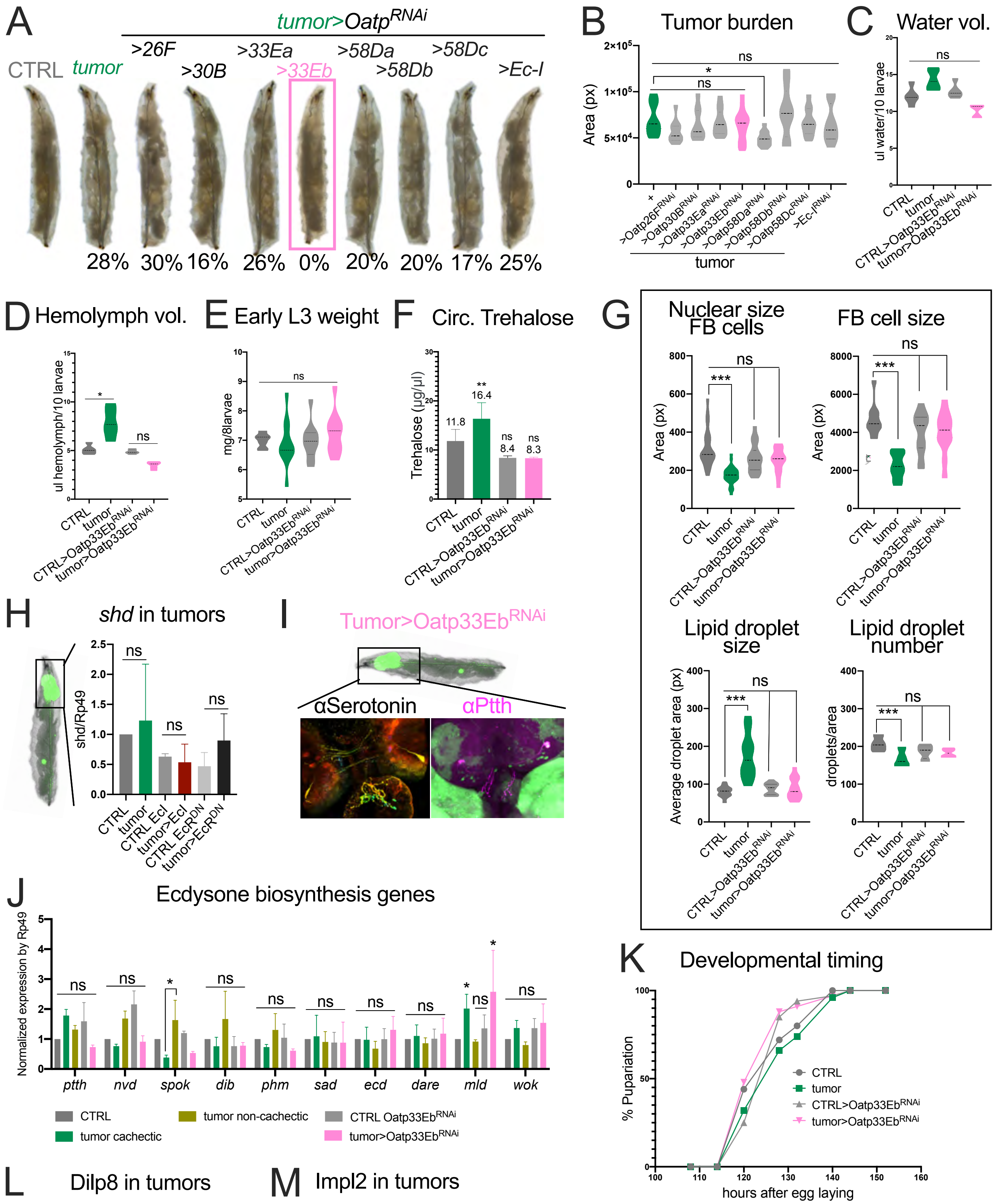


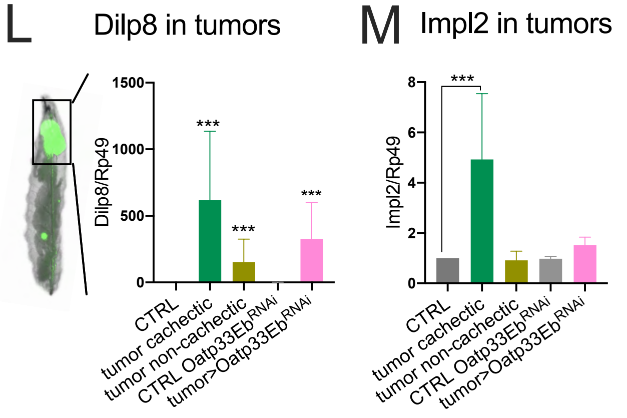


**N** EcR::Usp Binding sites in *Oatp33Eb* gene

| *Oatp33E sequence* | *Putative EcR::Usp BS* |
|---|---|
| Promoter (2000pb) | 13 |
| Promoter (5000pb) | 28 |
| 5'UTR+CDS+3'UTR+introns | 10 |

**Supplemental Figure 5. Supporting information for Figure 4.**

(A) Magnified larvae with the indicated genotypes. In all the cases, a percentage of tumor-bearing larvae display swelling defects except after the inhibition of the *Oatp33Eb* gene.

(B) Tumor sizes in larvae after the individual inhibition of each *Oatp* gene with an RNAi line (n=30 per genotype). The dotted line represents the mean.

(C,D) Water volume (C, n=50 for each genotype) and volumes of extractable hemolymph (D, n=50 for each genotype). The quantity of hemolymph is restored after *Oatp33Eb* inhibition in the tumor. The dotted line represents the mean. *p<0,05.

(E) Larval weight (n=11 groups of 8 larvae each one). No significant differences were observed.

(F) Concentration of Trehalose in the hemolymph from larvae with the indicated genotypes (mean +/- SEM, **p<0,01).

(G) Quantification of different parameters of fat body cells: Size of the nuclei (n=50), cell size (n=50), lipid droplet size (n=50) and lipid droplet number (5 groups in different areas). The dotted line represents the mean. ***p<0,001.

(H,J) mRNA levels of *shd* and genes involved in ecdysone synthesis in different types of tumors measured by RT-PCR. The results shown are mean +/- SEM. *mld* is upregulated in cachectic tumors and *tumors>Oatp33Eb*$^{RNAi}$, indicating that its expression does not reflect the cachectic state. *p<0,05, **p<0,01, ***p<0,001.

(I) Serotonin and Ptth stainings in the brain and prothoracic gland (PG) of *tumors>Oatp33Eb*$^{RNAi}$–bearing larvae.

(K) Effects on the developmental timing after tumor formation at the indicated time points. Pupariation is slightly advanced in *CTRL>Oatp33Eb*$^{RNAi}$ and *tumors>Oatp33Eb*$^{RNAi}$ animals.

(L,M) mRNA levels of *Dilp8* and *Impl2* from different types of tumors measured by RT-PCR. The results shown are mean +/- SEM. *p<0,05, ***p<0,001.

(N) Putative EcR::Usp binding sites in the *Oatp33Eb* promoter region.

**Genotypes**

**Fig. 1, Fig. 2, Fig. 3, Suppl. Fig. 1, Suppl. Fig. 2, Suppl. Fig. 3.**

CTRL: *eyFLP; +/CyO*$_{Gal80}^{TS}$*; act>y*$^{+}$*>Gal4, UAS-GFP/+*

Tumor: *eyFLP; UAS-Ras*$^{v12}$*, UAS-dlg*$^{RNAi}$*/+; act>y*$^{+}$*>Gal4, UAS-GFP/+* (*both cachectic and non-cachectic tumors have the same genotype)

CTRL>Ec-I: *eyFLP; UAS-Ec-I/CyO*$_{Gal80}^{TS}$*; act>y*$^{+}$*>Gal4, UAS-GFP/+*

Tumor>Ec-I: *eyFLP; UAS-Ras*$^{v12}$*, UAS-dlg*$^{RNAi}$*/UAS-Ec-I; act>y*$^{+}$*>Gal4, UAS-GFP/+*

CTRL>EcR$^{DN}$: *eyFLP; UAS-EcR*$^{DN}$*/CyO*$_{Gal80}^{TS}$*; act>y*$^{+}$*>Gal4, UAS-GFP/+*

Tumor>EcR$^{DN}$: *eyFLP; UAS-Ras*$^{v12}$*, UAS-dlg*$^{RNAi}$*/UAS-EcR*$^{DN}$*; act>y*$^{+}$*>Gal4, UAS-GFP/+*

CTRL>shd$^{RNAi}$: *eyFLP; UAS-shd*$^{RNAi}$*/CyO*$_{Gal80}^{TS}$*; act>y*$^{+}$*>Gal4, UAS-GFP/+*

Tumor>shd$^{RNAi}$: *eyFLP; UAS-Ras*$^{v12}$*, UAS-dlg*$^{RNAi}$*/UAS-shd*$^{RNAi}$*; act>y*$^{+}$*>Gal4, UAS-GFP/+*

CTRL>Cyp18a1: *eyFLP; UAS-Cyp18a1/CyO*$_{Gal80}^{TS}$*; act>y*$^{+}$*>Gal4, UAS-GFP/+*

Tumor>Cyp18a1: *eyFLP; UAS-Ras*$^{v12}$*, UAS-dlg*$^{RNAi}$*/UAS-Cyp18a1; act>y*$^{+}$*>Gal4, UAS-GFP/+*

**Sup. Fig. 1F: tGPH reporter activity.**

CTRL: *eyFLP; +/CyO*$_{Gal80}^{TS}$*; act>y*$^{+}$*>Gal4, UAS-GFP/tGPH*

Tumor: *eyFLP; UAS-Ras*$^{v12}$*, UAS-dlg*$^{RNAi}$*/+; act>y*$^{+}$*>Gal4, UAS-GFP/tGPH*

CTRL>Ec-I: *eyFLP; UAS-Ec-I/CyO*$_{Gal80}^{TS}$*; act>y*$^{+}$*>Gal4, UAS-GFP/tGPH*

Tumor>Ec-I: *eyFLP; UAS-Ras*$^{v12}$*, UAS-dlg*$^{RNAi}$*/UAS-Ec-I; act>y*$^{+}$*>Gal4, UAS-GFP/tGPH*

CTRL>EcR$^{DN}$: *eyFLP; UAS-EcR*$^{DN}$*/CyO*$_{Gal80}^{TS}$*; act>y*$^{+}$*>Gal4, UAS-GFP/tGPH*

Tumor>EcR$^{DN}$: *eyFLP; UAS-Ras*$^{v12}$*, UAS-dlg*$^{RNAi}$*/UAS-EcR*$^{DN}$*; act>y*$^{+}$*>Gal4, UAS-GFP/tGPH*

**Figure 2 A,B: EcRE_GFP.**

CTRL: *eyFLP; +/CyO*$_{Gal80}^{TS}$*; act>y*$^{+}$*>Gal4, UAS-GFP/EcRE_GFP*

Tumor: *eyFLP; UAS-Ras*$^{v12}$*, UAS-dlg*$^{RNAi}$*/+; act>y*$^{+}$*>Gal4, UAS-GFP/EcRE_GFP*

CTRL>Ec-I: *eyFLP; UAS-Ec-I/CyO*$_{Gal80}^{TS}$*; act>y*$^{+}$*>Gal4, UAS-GFP/EcRE_GFP*

Tumor>Ec-I: *eyFLP; UAS-Ras*$^{v12}$*, UAS-dlg*$^{RNAi}$*/UAS-Ec-I; act>y*$^{+}$*>Gal4, UAS-GFP/EcRE_GFP*

CTRL>EcR$^{DN}$: *eyFLP; UAS-EcR*$^{DN}$*/CyO*$_{Gal80}^{TS}$*; act>y*$^{+}$*>Gal4, UAS-GFP/EcRE_GFP*

Tumor>EcR$^{DN}$: *eyFLP; UAS-Ras*$^{v12}$*, UAS-dlg*$^{RNAi}$*/UAS-EcR*$^{DN}$*; act>y*$^{+}$*>Gal4, UAS-GFP/EcRE_GFP*

**Fig. 4, Suppl. Fig. 4.**

CTRL: *eyFLP; +/CyO*$_{Gal80}^{TS}$*; act>y*$^{+}$*>Gal4, UAS-GFP/+*

Tumor: *eyFLP; UAS-Ras*$^{v12}$*, UAS-dlg*$^{RNAi}$*/+; act>y*$^{+}$*>Gal4, UAS-GFP/+*

CTRL>Ec-I$^{RNAi}$: *eyFLP/UAS-Ec-I*$^{RNAi}$*; +/CyO*$_{Gal80}^{TS}$*; act>y*$^{+}$*>Gal4, UAS-GFP/+*

Tumor>Ec-I$^{RNAi}$: *eyFLP/UAS-Ec-I*$^{RNAi}$*; UAS-Ras*$^{v12}$*, UAS-dlg*$^{RNAi}$*/+; act>y*$^{+}$*>Gal4, UAS-GFP/+*

CTRL>Oatp26F$^{RNAi}$: *eyFLP; +/CyO*$_{Gal80}^{TS}$*; act>y*$^{+}$*>Gal4, UAS-GFP/ UAS-Oatp26F*$^{RNAi}$

Tumor>Oatp26F$^{RNAi}$: *eyFLP; UAS-Ras*$^{v12}$*, UAS-dlg*$^{RNAi}$*/+; act>y*$^{+}$*>Gal4, UAS-GFP/ UAS-Oatp26F*$^{RNAi}$

CTRL>Oatp30b$^{RNAi}$: *eyFLP; UAS-Oatp30b*$^{RNAi}$*/CyO*$_{Gal80}^{TS}$*; act>y*$^{+}$*>Gal4, UAS-GFP/+*

Tumor>Oatp30b$^{RNAi}$: *eyFLP; UAS-Ras*$^{v12}$*, UAS-dlg*$^{RNAi}$*/ UAS-Oatp30b*$^{RNAi}$*;act>y*$^{+}$*>Gal4, UAS-GFP/+*

CTRL>Oatp33Ea$^{RNAi}$: *eyFLP; UAS-Oatp33Ea*$^{RNAi}$*/CyO*$_{Gal80}^{TS}$*; act>y*$^{+}$*>Gal4, UAS-GFP/+*

Tumor>Oatp33Ea$^{RNAi}$:*eyFLP; UAS-Ras*$^{v12}$*, UAS-dlg*$^{RNAi}$*/ UAS-Oatp33Ea*$^{RNAi}$*;act>y*$^{+}$*>Gal4, UAS-GFP/+*

CTRL>Oatp33Eb$^{RNAi}$: *eyFLP; UAS-Oatp33Eb*$^{RNAi}$*/CyO*$_{Gal80}^{TS}$*; act>y*$^{+}$*>Gal4, UAS-GFP/+*

Tumor>Oatp33Eb$^{RNAi}$:*eyFLP; UAS-Ras*$^{v12}$*, UAS-dlg*$^{RNAi}$*/ UAS-Oatp33Eb*$^{RNAi}$*;act>y*$^{+}$*>Gal4, UAS-GFP/+*

CTRL>Oatp58Da$^{RNAi}$: *eyFLP; +/CyO*$_{Gal80}^{TS}$*; act>y*$^{+}$*>Gal4, UAS-GFP/UAS-Oatp58Da*$^{RNAi}$

Tumor>Oatp58Da$^{RNAi}$: *eyFLP; UAS-Ras*$^{v12}$*, UAS-dlg*$^{RNAi}$*/+; act>y*$^{+}$*>Gal4, UAS-GFP/ UAS-Oatp58Da*$^{RNAi}$

CTRL>Oatp58Db$^{RNAi}$: *eyFLP; UAS-Oatp58Db*$^{RNAi}$*/CyO*$_{Gal80}^{TS}$*; act>y*$^{+}$*>Gal4, UAS-GFP/+*

Tumor>Oatp58Db$^{RNAi}$: *eyFLP; UAS-Ras*$^{v12}$*, UAS-dlg*$^{RNAi}$*/ UAS-Oatp58Db*$^{RNAi}$*;act>y*$^{+}$*>Gal4, UAS-GFP/+*

CTRL>Oatp58Dc$^{RNAi}$: *eyFLP; +/CyO*$_{Gal80}^{TS}$*; act>y*$^{+}$*>Gal4, UAS-GFP/UAS-Oatp58Dc*$^{RNAi}$

Tumor>Oatp58Dc$^{RNAi}$: *eyFLP; UAS-Ras*$^{v12}$*, UAS-dlg*$^{RNAi}$*/+;act>y*$^{+}$*>Gal4, UAS-GFP/UAS-Oatp58Dc*$^{RNAi}$
